# The infiltration pattern of microenvironmental cells and different immune escape mechanisms in colorectal cancer

**DOI:** 10.1101/2022.02.16.480691

**Authors:** Yihao Mao, Qingyang Feng, Wenju Chang, Yang Lv, Yuqiu Xu, Jiang Chang, Peng Zheng, Shanchao Yu, Zhiyuan Zhang, Zhiqiang Li, Qi Lin, Wentao Tang, Dexiang Zhu, Meiling Ji, Li Ren, Ye Wei, Guodong He, Jianmin Xu

## Abstract

**Background:** The tumour microenvironment (TME) plays a crucial role in tumour progression and metastasis. However, the infiltration pattern of TME cells in CRC samples and the immune escape mechanism underneath have not been studied sufficiently.

**Methods:** Transcriptomic data from public datasets were retrieved online. In total, 1802 samples from the microarray dataset and 619 samples from the TCGA dataset were enrolled. The ssGSEA algorithm and unsupervised clustering were used for TME cells infiltration speculation and infiltration pattern recognition.

**Results:** CRC samples can be classified into three distinct TME cell subtypes. Subtype 1, the immune-active subtype, was characterised by high infiltration of activated adaptive immune cells. Subtype 2, the immune-desert subtype, featured high tumour purity and low infiltration of immune and stromal cells. And subtype 3, stroma-rich subtype, had high infiltration of stromal cells. The stroma-rich subtype conferred a significantly worse prognosis. Three subtypes had different immune escape mechanisms. The immune-active subtype has the highest immune checkpoint expression level. In comparison, the immune-desert subtype had the lowest immunogenicity and the defective antigen presentation. And the stroma-rich subtype lacked activated immune cells.

**Conclusions:** Distinct TME cell subtypes and immune escape mechanisms may provide inspiration and direction for further researches on CRC immunotherapy.

## 1 Introduction

Colorectal cancer (CRC) is estimated to be the third most commonly diagnosed and cause of death in both men and women in the USA in 2021[1]. Tumour microenvironment (TME), defined as the surrounding environment of a tumour, comprises tumour cells, immune cells, stromal cells, blood vessels, and other mesenchymal cells. TME plays a crucial role in tumour progression and metastasis[2].

Researchers have focused on CRC TME from multiple perspectives. Multiple TME cells were found to have prognostic value and predict therapeutic benefit [3, 4]. As a classic example, Immunoscore, which quantifies the in situ T cell infiltration in CRC tissue, was validated to be a valuable prognostic factor[5] and was recommended in the European Society for Medical Oncology (ESMO) guidelines[6]. However, numerous studies only focused on one or few types of TME cells. Moreover, most of them were based on pathology level with a limited sample size. Confusing, even contradictory results were gotten. As CRC was a highly heterogeneous cancer, we assumed that the same type of TME cell might play different roles in different CRC tissues.

Recently, emerging papers explored the CRC TME using the bioinformatic based method, such as CIBERSORT[7], MCPcounter[8], TIMER[9], et al. However, because of the limitation of the above methods, like only relative infiltration results provided or limited predictable cell types, the landscape of TME cell infiltration pattern in CRC samples has not been systematically clarified to data, nor did the immune escape mechanism underneath elucidated.

In this study, we comprehensively explored the TME cell infiltration pattern in two large independent cohorts. CRC samples can be divided into three subtypes with distinct TME cell infiltration patterns and their immune escape mechanism underneath. Also, the different clinical, prognostic, and genomic characteristics were described.

## 2 Materials and methods

### 2.1 Study population

Public datasets selection criterion was described as follows: (1) transcriptomic data, including microarray data using Affymetrix HG-U133A (GEO accession number GPL96) or HG-U133 Plus 2.0 (GEO accession number GPL570) platforms or RNA-Seq data, were available; (2) the basic clinicopathological information, including AJCC/UICC TNM stage and survival information (overall survival (OS) or disease-free survival (DFS)) was available; (3) the sample size was larger than 50. After systematically searching, GSE14333[10], GSE17538[11], GSE33113[12], GSE37892[13], GSE38832[14], GSE39084[15], GSE39582[16], KFSYSCC, TCGA-COAD and TCGA-READ datasets[17] were included in our research. Finally, 1802 samples from the microarray dataset and 619 samples from the TCGA dataset were enrolled for subsequent analysis. The composition of each dataset was illustrated in Fig. S1. The microarray dataset was utilised to discover the infiltration pattern of TME cells and perform survival-related analysis. Subsequent pattern validation and mechanism exploration were conducted in the TCGA dataset. Due to the relatively shorter follow-up time of the TCGA dataset (median, 21.9 months), survival-related analysis was only performed in the microarray dataset.

### 2.2 Data acquisition and preprocessing

The transcriptome and clinical data of GSE14333, GSE17538, GSE33113, GSE37892, GSE38832, GSE39084, and GSE39582 were downloaded from Gene Expression Omnibus (GEO) repository (https://www.ncbi.nlm.nih.gov/gds/). The expression matrix and clinical information of KFSYSCC were acquired from the supplementary material of a previous publication[18]. The RNA-Seq (RSEM normalised), copy number variation (CNV), mutation, images of pathology slides, and clinical data of TCGA-COAD and TCGA-READ datasets were downloaded from Genomic Data Commons (GDC, https://portal.gdc.cancer.gov/) in Dec. 2019.

We used the voom function (in R package “limma” [19]) to transform RSEM normalised TCGA RNA-Seq expression matrix. The ComBat algorithm (“ComBat” function in R package “sva” [20]) was applied to calibrate the heterogeneity among different datasets in microarray and TCGA datasets, respectively. The PCA plots illustrating the effect of calibration were shown in Fig. S2.

### 2.3 Microenvironmental cell infiltration speculation and validation

We used the ssGSEA algorithm (“gsva” function in R package “GSVA” [21]) for infiltration speculation of 31 microenvironmental cells based on transcriptomic data as previously described[22, 23]. Gene signature used in our study was mainly based on signature described in CHAROENTONG P et al. [24], XIAO Y et al.[22] (which was a modification of LM22 signature of CIBERSORT) and MCPcounter[25]. The detailed gene signature was described in Table S1.

To validate the accuracy of our methods, we compared our calculation results with the estimation of the cell abundances by CIBERSORT and MCPcounter (Table S2 and S3). The comparison results showed high accordance between methods.late. These format

### 2.4 Unsupervised clustering based on TME cell infiltration pattern

For TME cell infiltration pattern discovery and validation, a nonnegative matrix factorisation algorithm (NMF) (“nmf” function in R package “NMF” [26]) was performed for unsupervised clustering on the scaled ssGSEA results. We used the “nmfEstimateRank” function (30 runs) to choose the optimal number of clusters based on the Cophenetic correlation coefficient changing. The highest clustering number before the Cophenetic correlation coefficient dropping most was selected as the optimal clustering number.

### 2.5 Slide Image analysis

Hematoxylin and eosin (HE) stained diagnostic slides of the TCGA dataset were retrieved from GDC Data Portal. The infiltration of tumour-infiltrating lymphocytes (TILs) was recorded as the mean number of lymphocytes in tumour tissue from three random high power fields (HPF, 200×). Furthermore, the infiltration of stromal cells was evaluated as the mean proportion of stromal cells in tumour tissue from three randomised HPFs. Tumour necrosis in HE slides was characterised by degraded tumour cells, presented as amorphous coagulum with nuclear debris (Fig 3G)[27]. Pathologists annotated the necrosis area in each diagnostic slide and the proportion of necrosis was calculated by ImageJ (US National Institutes of Health, Bethesda, MD, USA). The above results were all assessed by two independent pathologists who were blinded to the clinical data and the results were averaged.

**Figure 1.**
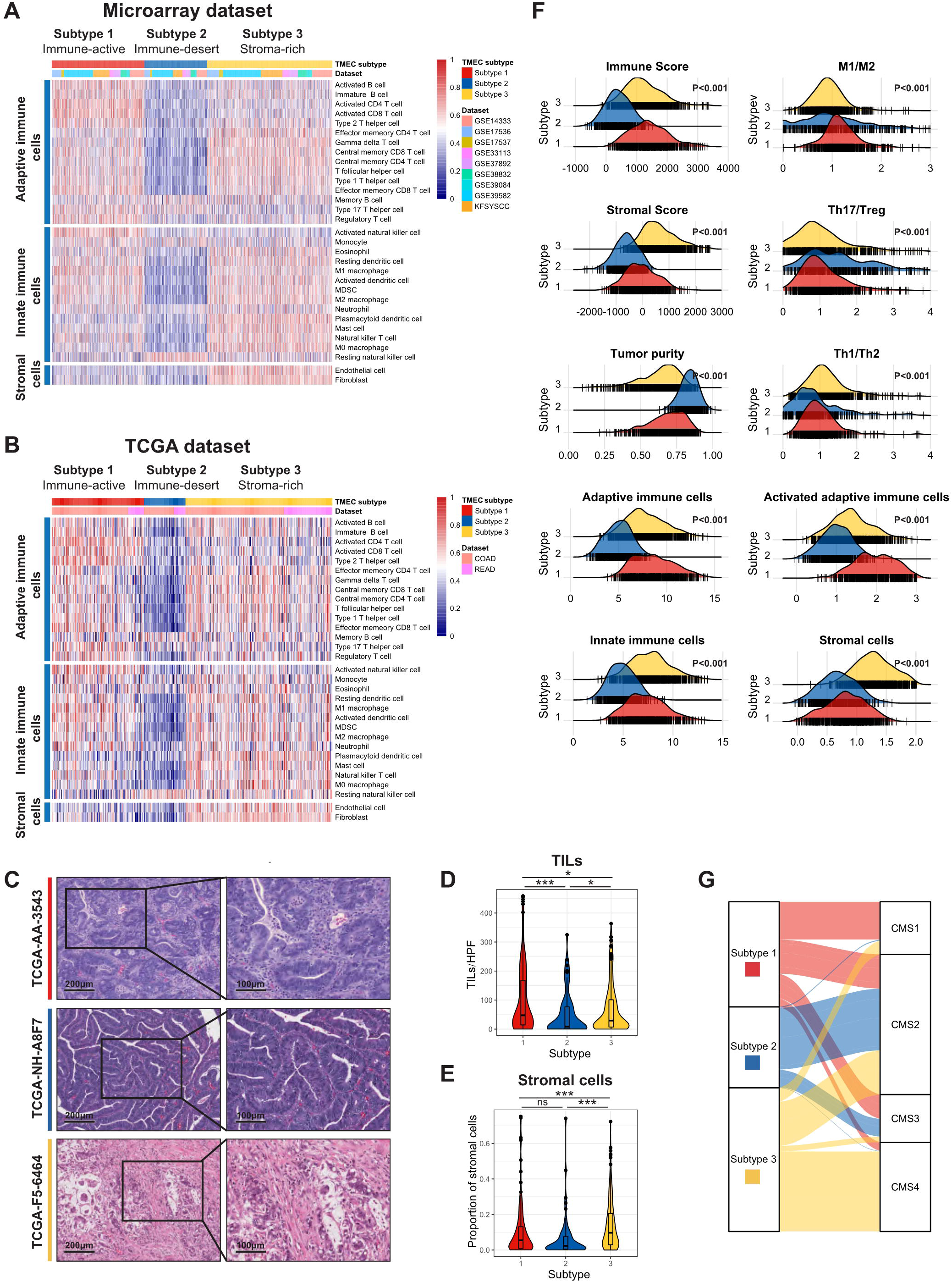
The landscape of TME cell infiltration pattern in CRC. (A-B) Consensus heatmap illustrating the infiltration of 31 TME cells in the microarray dataset (A) and the TCGA dataset (B). Both of them were divided into three subtypes. (C) The representative pathology presentation of each subtype in the TCGA dataset (red, subtype 1; blue, subtype 2; yellow, subtype3; from top to bottom). (D) The number of TILs among three subtypes per high power field (HPF, 200×) in the TCGA dataset. (E) The proportion of stromal cells among three subtypes per high power field (HPF, 200×) in the TCGA dataset. (F) The distribution of Immune Score, Stromal Score, tumour purity, macrophage M1/M2 ratio, Th17/Treg ratio, Th1/Th2 ratio, adaptive immune cells, activated adaptive immune cells, innate immune cells, and stromal cells among subtypes in the microarray dataset. (G) Sankey plot illustrating the distribution of CMS subtypes and TMEC subtypes in the microarray dataset.

**Figure 2.**
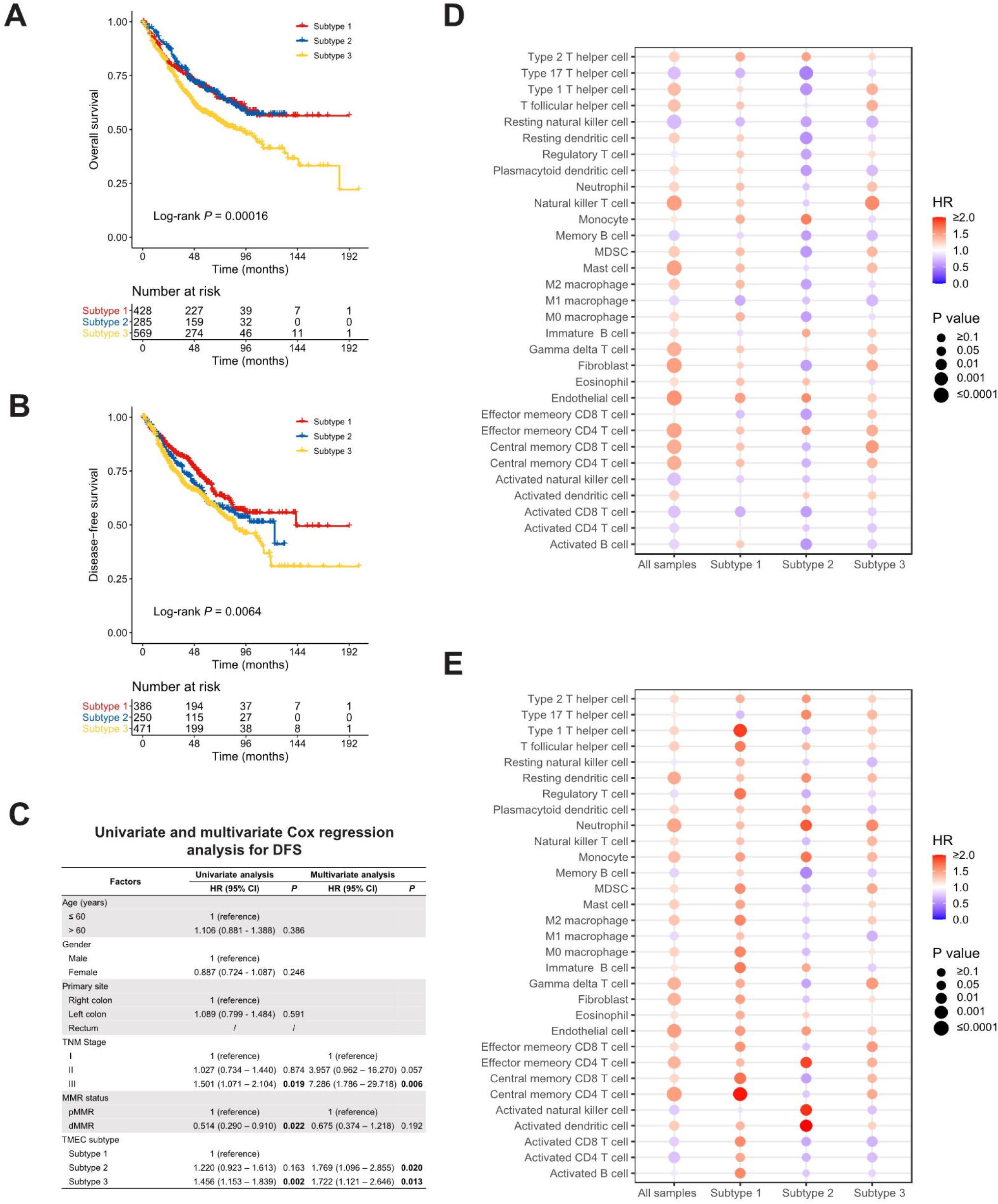
(A-B) Kaplan-Meier analysis of OS (A) and DFS (B) in the microarray dataset. (C) Univariate and multivariate Cox regression of clinicopathological factors and TMEC subtypes for DFS in the microarray dataset. (D-E) The prognostic value of different TME cells in all samples and each subtype in the microarray dataset for OS (D) and DFS (E).

**Figure 3.**
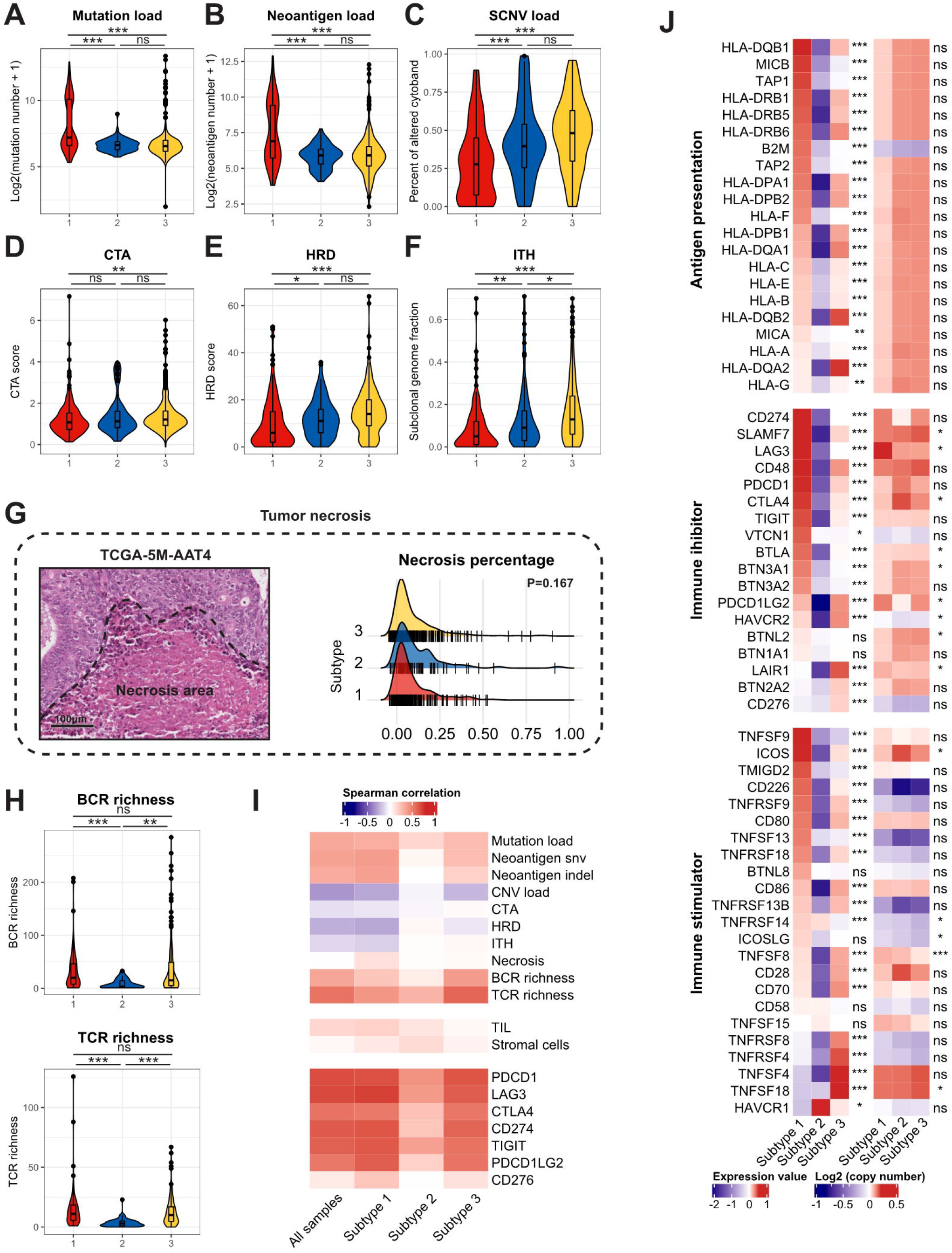
(A-F) The mutation load (A), neoantigen load (B), SCNV load (C), CTA (D), HRD (E) and ITH (F) among TMEC subtypes. (G) The representative pathology image of tumor necrosis (left) and the distribution of necrosis percentage of each subtype. (H) The BCR and TCR richness of different subtypes. (I) The correlation between immune cytolytic activity and tumour immunogenicity factors, TILs, stromal cells, and immune checkpoint molecules. (J) The relative expression levels to the median value and mean log2(copy number) value of immune inhibitors and immune stimulators among three subtypes. ***: P < 0.001; **: 0.001 < P < 0.01; *: 0.01 < P < 0.05; ns: P > 0.05.

### 2.6 Genomic analysis

TCGA level 3 mutation data were downloaded from GDC Data Portal. The mutation load was defined as log2(non-silent mutation number + 1). The neoantigen data was downloaded from The Cancer Immunome Atlas (TCIA, https://tcia.at/)[28]. The cancer-testis antigen (CTA) scores, homologous recombination deficiency (HRD) scores were downloaded from the supplementary material of a previous publication[23]. The intratumor heterogeneity (ITH), defined as the subclonal genome fraction measured by ABSOLUTE, was retrieved from an earlier publication[23]. The GISTIC2.0 results of In silico Admixture Removal (ISAR) calibrated SCNV data (minus germline CNV) were obtained from a previous publication[23]. The GISTIC2.0 thresholded result of 2 or −2 was defined as deep CN alterations called amplifications and depletions. While the result of 1 or −1 was defined as shallow CN alterations, which were called gains and losses, respectively. The SCNV load was calculated as the percentage of altered cytoband in each sample. T cell & B cell receptor (TCR&BCR) diversity scores (Shannon Entropy, Evenness, and Richness) were obtained from a previous publication[23]. Immune cytolytic activity score (CYT), which highly correlated with CD8+ T cell activation, was defined as the log-average (geometric mean) of GZMA and PRF1 expression as previously described[29].

### 2.7 Mutations and CNVs comparison between subtypes

Mutations and CNVs were compared between every two subtypes. To adjust the difference of mutation burden between subtypes, we adopted the method based on logistic regression as described in a previous publication[22]. Specifically, the subtype was modelled as a logistic function as subtype ~ mutation burden + gene mutation status. If the P-value of a specific gene mutation status adjusted for mutation burden less than 0.05, such gene was defined as differentially mutated between subtypes. Notably, gene mutations with an overall mutation rate of less than 2.5% were excluded from the comparison. CNVs were compared similarly. Gains/amplifications and losses/deletions were compared between every two subtypes separately. For gains/amplifications comparison, losses/deletions events were set to zero. The subtype was modelled as a logistic function as subtype ~ CNV burden + gene gain/amplification status. If the P-value of a specific gene gain/amplifications adjusted for CNV burden less than 0.05, such gain/amplification was defined as differentially distributed between subtypes. The losses/deletions comparisons were performed similarly. The above calculations were performed using the “glm” function in R.

### 2.8 Statistical analysis

All statistical analyses were performed in R software, version 3.6.1 (The R Foundation for Statistical Computing, http://www.rproject.org/). Continuous and ordered categorical variables were compared using Student's t-tests or Kruskal–Wallis test with post hoc pairwise Dunn's-tests. Unordered categorical variables were compared by Pearson χ^2^ test or Fisher exact test. Spearman correlation analyses were employed for evaluating correlations between continuous variables. Survival analyses were performed using Kaplan–Meier analyses and survival differences between groups were compared by log-rank tests. Prognostic factors were identified by univariate and multivariate Cox regression analyses. Factors with univariate regression P<0.1 were further enrolled in multivariate analyses. A two-sided P<0.05 was considered statistically significant unless additionally stated.

## 3 Results

### 3.1 Microenvironmental cell infiltration pattern in CRC

We used the ssGSEA algorithm for estimating the absolute infiltration abundance of 31 different TME cells in 1802 samples of the microarray dataset and 619 samples of the TCGA dataset. After normalisation, a nonnegative matrix factorisation (NMF) algorithm, an unsupervised clustering method, was applied to classify CRC samples into different subtypes based on TME cell infiltration. The optimal cluster number of 3 was selected by the Cophenetic correlation coefficient in both datasets (Figs. S3 and S4). Thus, we classified 1802 samples into three distinct TME cell (TMEC) subtypes (Fig. 1A). Subtype 1, named as immune-active subtype (marked by red), was characterised by high infiltration of adaptive and innate immune cells, especially activated adaptive immune cells, such as activated CD4 T cells, activated CD8 T cells and activated B cells, and low infiltration of stromal cells. Subtype 2, named as immune-desert subtype (marked by blue), featured low infiltration of most immune and stromal cells. And subtype 3, named as stroma-rich subtype (marked by yellow), had high infiltration of adaptive and innate immune cells, as well as stromal cells but low infiltration of activated adaptive immune cells compared with subtype 1. The above infiltration pattern was also validated in the TCGA dataset (Fig. 1B).

To further validate the above subtypes, we evaluated the pathological presentation of three distinct subtypes in diagnostic slides of 499 TCGA samples. The representative pathology features of each subtype were illustrated in Fig. 1C. Tumour tissue in subtype 1 featured median stromal proportion with high tumor infiltrating leukocytes (TILs) infiltration. Subtype 2 had high tumour purity and low TME cell infiltration. And mesenchymal cells infiltrated extensively in subtype 3. Then we evaluated the TILs and stromal cell infiltration pathologically in TCGA diagnostic slides. TILs infiltrated most in subtype 1 and least in subtype 2 (median, 47 vs. 8 vs. 29, P<0.001), while subtype 3 had significantly more stromal cell proportion (median, 0.047 vs. 0.037 vs. 0.098, P<0.001).

Further, we explored other characteristics of each subtype. We used the ESTIMATE algorithm to calculate Immune Score, Stromal Score, and tumour purity for each sample. As shown in Fig. 1F, Subtype 1 had the highest Immune Score, median Stromal Score, and relatively low tumour purity. And subtype 2 featured the lowest Immune Score and Stromal Score but the highest tumour purity. Subtype 3 had a median Immune Score, highest Stromal Score, and relatively low tumour purity (all P<0.001). Also, The macrophage M1/M2 ratio was the highest in subtype 1 (P<0.001), while Th17/Treg ratio was the highest in subtype 2 (P<0.001). Samples in subtype 3 own the highest Th1/Th2 ratio (P<0.001) and the proportion of activated adaptive immune cells was the highest in subtype 1 (P<0.001). The above characteristics were all validated in the TCGA dataset (Fig. S6).

### 3.2 TMEC Subtypes and clinicopathological features

The correlation between TMEC subtypes and clinicopathological features of microarray and TCGA datasets was presented in Table S4 and S5, respectively. Patients in subtype 1 were relatively older (microarray dataset: mean±SD, 66.83±14.48 vs. 65.87±13.70 vs. 64.22±12.99, P = 0.005; TCGA dataset: mean±SD, 68.37±13.51 vs. 65.25±10.98 vs. 65.31±12.63, P = 0.019). Subtype 3 had more samples with advanced stages (P<0.001 in both datasets). Subtype 1 was associated with more right colon cancer (P<0.001 in both datasets). Also, the dMMR/MSI-H samples were significantly enriched in subtype 1 (P<0.001 in both datasets). Moreover, we compared the distribution of Consensus Molecular Subgroups (CMS)[30] and TMEC subtypes, illustrated as Sankey plots in Fig. 1G and Fig. S7. Subtype 1 was mainly composed of CMS1 (MSI immune), CMS2 (Canonical), and CMS3 (Metabolic) samples. The majority of subtype 2 samples were from CMS2. And samples of CMS2 and CMS4 (Mesenchymal) constituted most of subtype 3. The above features were presented in both microarray and TCGA datasets.

### 3.3 TMEC Subtypes and CRC prognosis

TMEC subtypes also plays an important role in CRC prognosis. The analyses related to overall survival (OS) were performed in 1281 samples from GSE17538/GSE38832/GSE39084/GSE39582/KFSYSCC with OS data available, while the analyses related to disease-free survival (DFS) were performed in 1107 samples from GSE14333/GSE17538/GSE33113/GSE38832/GSE39084/GSE39582 with DFS data available. Subtype 3 conferred significantly worse OS and DFS, while subtype 1 and subtype 2 had similar prognoses (Fig. 2A and 2B). Next, we conducted univariate and multivariate Cox regression of clinicopathological factors and TMEC subtypes for OS and DFS (Table S6 and Fig. 2C, respectively). TMEC SUBTYPE of subtype 3 was a significant prognostic indicator in univariate analyses of both OS (P<0.001, HR = 1.449, 95% CI = 1.174 −1.787) and DFS (P=0.002, HR=1.456, 95% CI = 1.153-1.839). Multivariate analyses showed subtype 2 and subtype 3 were independent prognostic factors of DFS (P=0.020, HR=1.769, 95% CI=1.096-2.855 and P=0.013, HR=1.722, 95% CI=1.121-2.646, respectively) (Fig. 2C) but not OS (Table S6).

Subgroup analyses of OS and DFS revealed that subtype 3 indicated impaired survival in most scenarios (Fig. S8). Furthermore, we analysed the prognostic value of different TME cells in all samples and each subtype in the microarray dataset for OS (Fig. 2D) and DFS (Fig. 2E). Interestingly, the same cell type may play converse prognostic roles in different TMEC subtypes. In subtype 2, the majority of immune and stromal cells presented as protective factors of survival, even immunoinhibitory cells like MDSCs and M2 macrophages, which indicated heterogeneous immune networks in different TMEC subtypes.

### 3.4 Possible immune escape mechanism of distinct TMEC subtypes

The distinct characteristics of TMEC subtypes made us wonder whether they had different immune escape mechanisms. Previous studies summarised vital factors leading to tumour immune escape as follows: (1) defective antigen presentation; (2) tolerance and immune deviation; (3) infiltration of immune-suppressive cells; (4) immune-suppressive mediators secretion [31, 32]. As multi-omics data was available for the TCGA dataset, we explored the possible immune escape mechanism of three TMEC subtypes.

#### 3.4.1 Defective antigen presentation

Defective antigen presentation was composed of at least two aspects: alteration of tumour immunogenicity and down-regulation of antigen presentation pathway. For tumour immunogenicity evaluation, subtype 1 had the highest mutation burden (Fig. 3A) and neoantigen load (Fig. 3B) (both P < 0.001), which could be easily speculated for the highest proportion of dMMR/MSI-H samples. While subtype 3 featured the highest SCNV load (Fig. 3C). The CTA and HRD scores of subtype 3 were higher than subtype 1 (P = 0.008 and P < 0.001, respectively) but not subtype 2 (both P > 0.05) (Fig. 3D, Fig. 3E, respectively). Also, subtype 3 had the highest ITH level (P < 0.001) (Fig 3F). The necrosis level between subtypes was not significantly differentiated (P = 0.167) (Fig. 3G). Subtype 2 showed the lowest BCR&TCR richness diversity (both P < 0.01) (Fig. 3H), which were positively correlated with cytolytic activity (Fig. 3I). As for antigen presentation pathway-related gene expression, subtype 1 had higher expression of most MHC-related genes, while the expression of subtype 2 was the lowest (all P < 0.01) (Fig. 3J). Overall, three subtypes all had impaired antigen presentation to some extent, but subtype 2 had the lowest immunogenicity and antigen presentation gene expression level.

#### 3.4.2 Tolerance and immune deviation

Tumours were also known to induce immune tolerance by upregulating immune inhibitors and downregulating immune stimulators. The relative expression levels and mean log2(copy number) value of immune inhibitors and immune stimulators were illustrated in Fig. 3J. Subtype 1 had the highest expression level of most immune inhibitors and part of immune stimulators, which indicated pro-tumour and anti-tumour immunity activation. The activation level of most above genes was the lowest in subtype 2. Subtype 3 highly expressed TNF/TNF-receptors. Some differentially expressed genes, such as SLAMF7, LAG3, CTLA4, BTLA, TNFSF8, et al. in two modules might attribute to SCNV. Correlation analyses found that immune cytolytic activity was positively associated with most immune checkpoint expression (Fig 3I). Generally, subtype 1 had a high expression of both immune stimulators and immune inhibitors, whose expression level was the lowest in subtype 2.

#### 3.4.3 Infiltration of immune-suppressive cells

Next, we compared the infiltration of each cell type between subtypes, illustration as volcano plots (Fig.4A). Comparing to subtype 2 and subtype 3, subtype 1 had significantly more activated CD4 and CD8 T cells infiltration. Subtype 2, as an immune-cold subtype, had the least adaptive and innate immune cells infiltration among three subtypes. For subtype 3, it had enriched fibroblasts and endothelial cells than the other two subtypes. Immune-suppressive cells, such as Tregs, MDSCs, M2 macrophages, were more abundant in subtype 1 and 3. Also, we used the gene signature from a high-quality publication[33] for exhausted T cells infiltration speculation and compared the infiltration of exhausted T cells in each subtype (Fig. 4B). Subtype 1 had a significantly higher exhausted T cell proportion than the other two subtypes (P < 0.001), as well as the ratio of exhausted T cells to CD8 T cells (P < 0.001).

**Figure 4.**
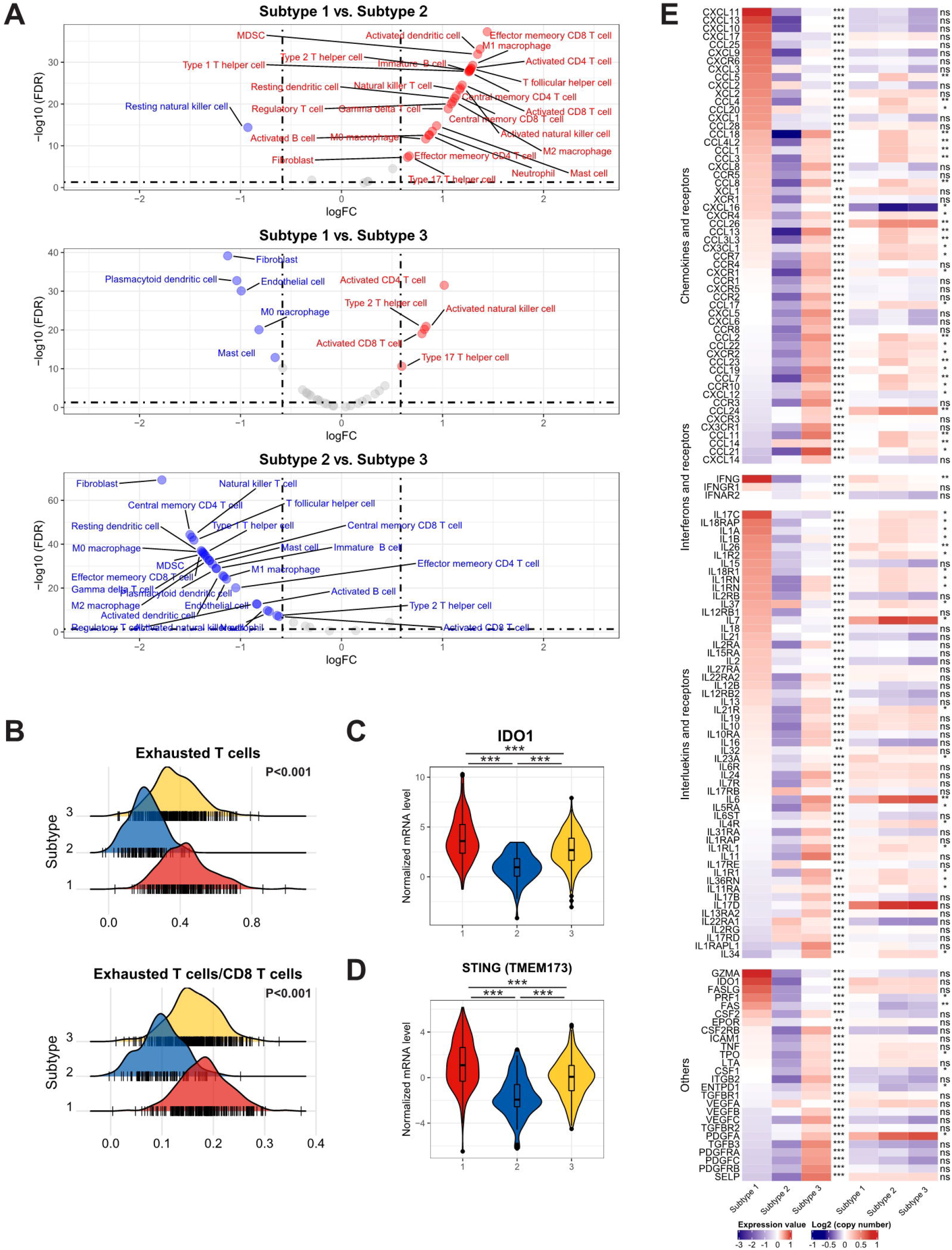
(A) The comparison of each cell type between subtypes. (B) The distribution of exhausted T cells infiltration and the ratio of exhausted T cells to CD8 T cells infiltration among each subtype. (C-D) The expression of IDO and STING among each subtype. (E) The relative expression levels to the median value and mean log2(copy number) value of chemokines, interleukins, other cytokines and their receptors among three subtypes. ***: P < 0.001; **: 0.001 < P < 0.01; *: 0.01 < P < 0.05; ns: P > 0.05.

#### 3.4.4 Immune suppressive mediators secretion

Furthermore, we investigated the fourth aspect of the immune escape mechanism. The relative expression levels to the median value and mean log2(copy number) value of chemokines and receptors, interferons and receptors, interleukins and receptors, and other cytokines in three subtypes were compared, and cytokines with P < 0.01 was illustrated in Fig. 4E. Most cytokines and receptors expression were negatively correlated between subtype 1 and subtype 3, and the expression level of subtype 2 was the lowest on most occasions. Subtype 1 expressed higher pro-immune chemokines like CXCL9, CXCL10, CXCL11, which could recruit effector T cells and NK cells[34]. The IFN-γ, which robustly stimulated anti-tumour immunity, was expressed higher in subtype 1. Pro-immunity IL-2 and IL-15 were accumulated in subtype 1. IL-34, CCL2, CCL-3, CXCL8, and some of their receptor CCR5, CXCR1, CXCR2, which promoted homeostasis of myeloid cells[35], were expressed extensively in subtype 3. Mesenchymal development-related PDGFs and PDGFRs[36] and cell adhesion-related SELP, ICAM1, ITGB2 were enriched in subtype 3. However, IDO, which contributed to peripheral tolerance, was significantly highly expressed in subtype 1 (P < 0.001) (Fig. 4C), indicating the existence of inhibitory cytokines in the inflamed microenvironment. Notably, subtype 2 expressed a significantly higher level of VEGFA. Moreover, STING, a crucial factor in the cGAS–Sting pathway, which contributes to the initiation of innate immunity and recognition of the tumour, is highly expressed in subtype 1 (P < 0.001) (Fig. 4D), suggesting the impaired immunity initiation in subtype 2 and 3.

### 3.5 TMEC subtype-specific genomic alterations

Other than the heterogeneous immune escape mechanism among the TMEC subtypes, we further explored the subtype-specific genomic alterations. First, we explored the activation of each subtype in a comprehensive cancer development-related pathways collection [37, 38]. As presented in Fig. 5A, TP53, NRF2, PI3K pathways were upregulated in subtype 1, while MYC, PI3K pathways were enriched in subtype 2. Hippo, NOTCH, Hedgehog, TGF-β, Wnt, RAS pathways were activated in subtype 3. As for other cancer development-related pathways, cell cycle, genomic repair, protein expression, metabolism, and immunity pathways were generally upregulated in subtype 1. The immunity and stromal pathways, which were highly enriched in subtype 3, were relatively suppressed in subtype 2.

**Figure 5.**
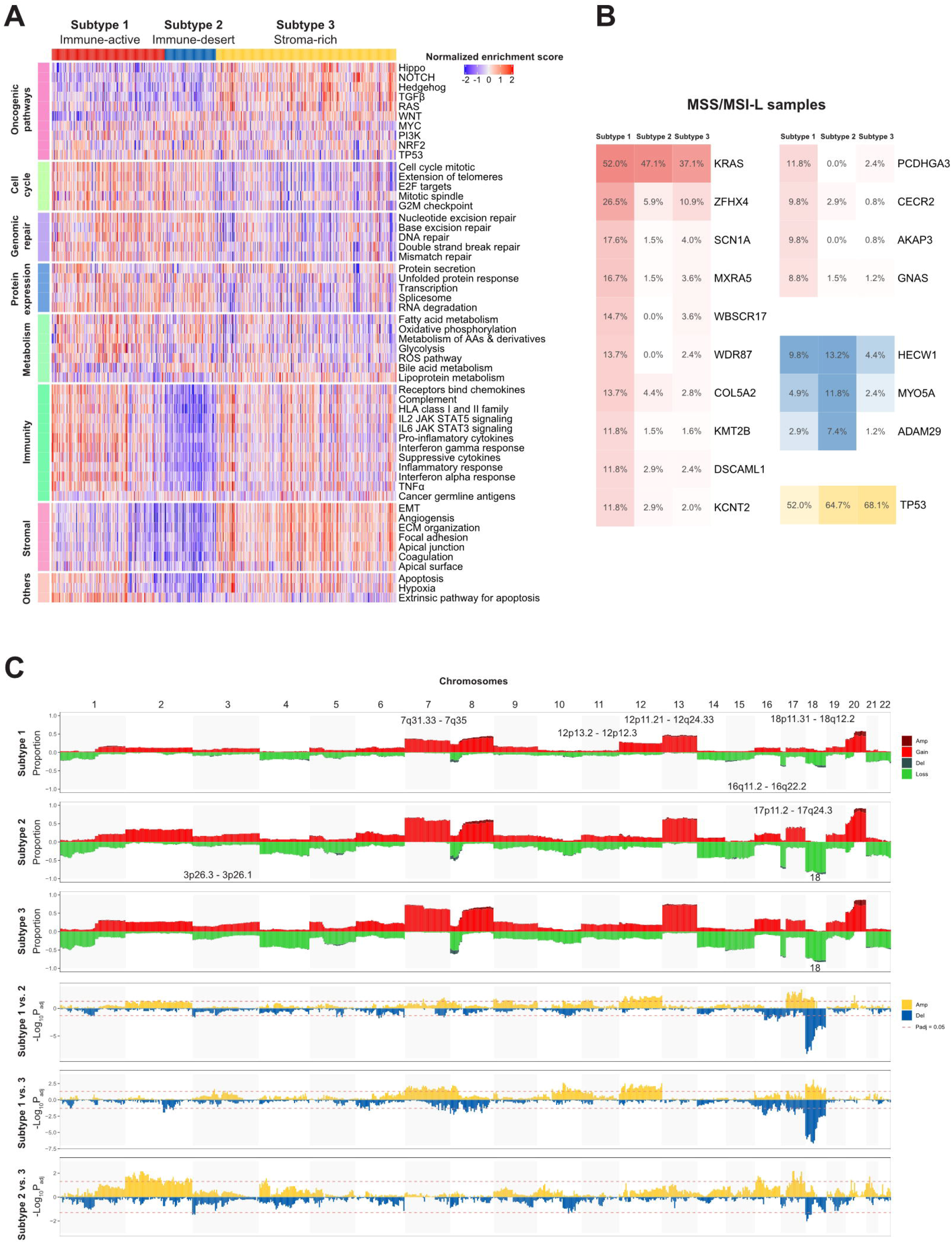
TMEC subtype-specific genomic alterations. (A) Heatmap illustrating activation of comprehensive cancer development-related pathways among three TMEC subtypes. (B) Subtype-specific gene mutation frequency among MSS/MSI-L samples in three TMEC subtypes. (C) Subtype-specific SCNVs of each TMEC subtype (top three plots) and comparison of SCNVs between every two subtypes.

Next, we compared the subtype-specific gene mutations. Due to the high percentage of MSI-H patients in subtype 1, it is easy to speculate that most mutations were enriched in subtype 1. However, most mutations in MSI-H patients were random and irregular. Therefore, we focused on the comparison among MSS/MSI-L patients. As shown in Fig. 5B, after adjusting for mutation burden, KRAS, ZFHX4, SCN1A, MXRA5, WBSCR17, WDR87, COL5A2, KMT2B, DSCAML1, KCNT2, PCDHGA3, CECR2, AKAP3, and GNAS were highly mutated in subtype 1. And the mutation frequency of HECW1, MYO5A, and ADAM29 was higher in subtype 2, While TP53 mutated more in subtype 3 (all adjusted P <0.05).

As the CNV load of subtype 1 was significantly lower than the other two subtypes, the proportion of alteration in most cytoband was the lowest among subtypes (Fig 5C). However, after adjusting for CNV burden, most cytobands in Chr12 were more frequently amplificated in subtype 1. Also, 18p11.31-18q12.2 amplification was enriched in subtype 1. Subtype 2 exhibited more frequent 17p11.2-17q24.3 gain/amplification. Moreover, chr18 loss/deletion was more frequent in subtype 2 and 3.

## 4 Discussion

The development of personalised cancer medicine has garnered the concern of CRC subtyping. The CRC subtyping system developed from pathological staging, mutation-based subtyping to omic-based subtyping[30]. However, the large-scale comprehensive presentation of CRC TME cell infiltration pattern and subsequent subtyping research was rare and insufficient. This study systematically showed the infiltration pattern of primary TME cells in two large independent cohorts, the microarray dataset and the TCGA dataset for cross-validation. The CRC samples can be mainly divided into three subtypes, immune-active, immune-desert, and stroma-rich. The major characteristics of each subtype were concluded in Fig. 6. The immune-active subtype features high activated adaptive immune cells infiltrate, more right colon, more dMMR/MSI-H, and more CMS1 samples. The immune-desert subtype was characterised by low infiltration of most TME cells, more left colon, and more CMS2. And the stroma-rich subtype presents the distinctiveness of high infiltration of both immune cells and stromal cells, advanced stages, poor prognosis, and more CMS4 samples. Similar TME subtypes can also be observed in ovarian cancer[39] and TNBC[22].

**Figure 6.**
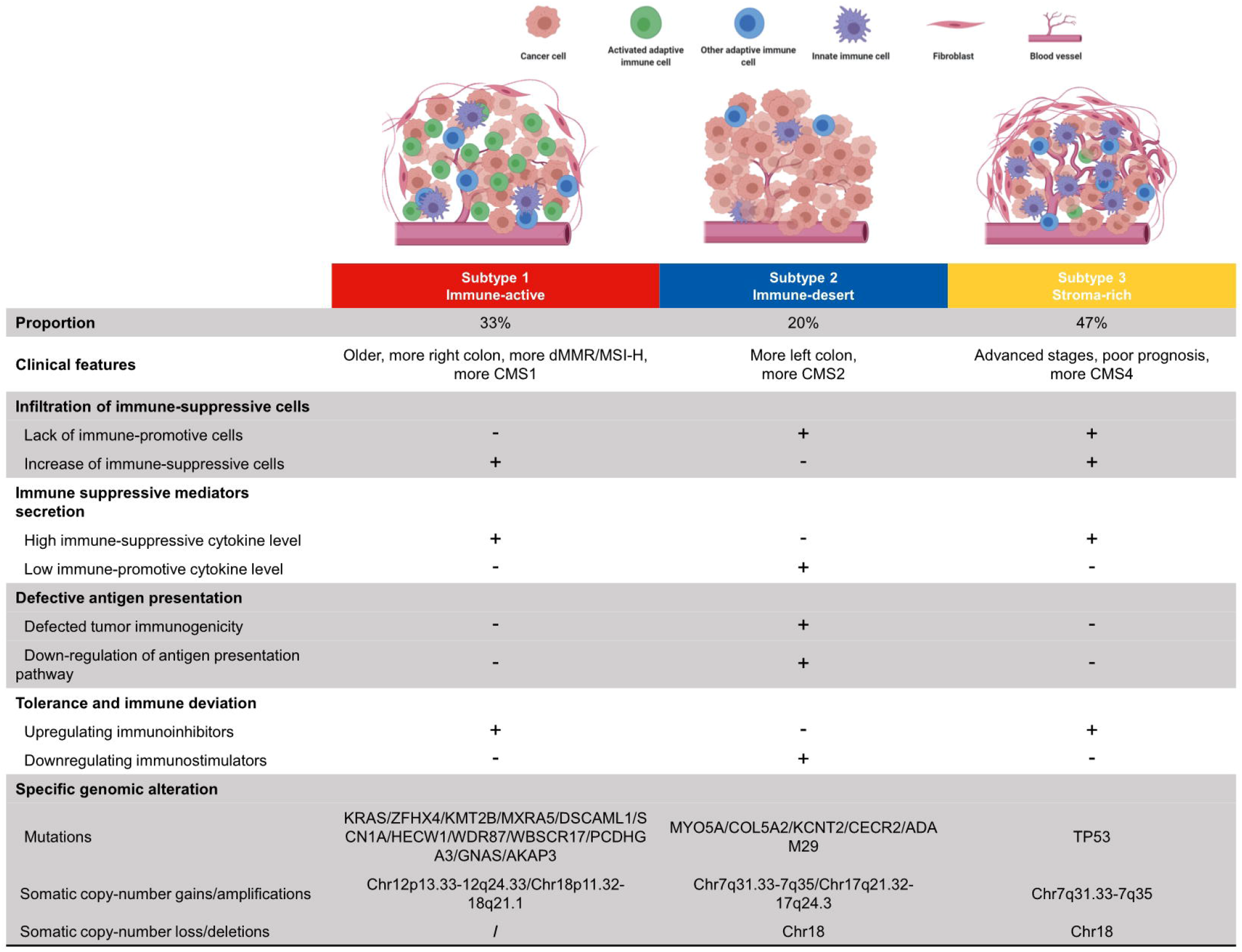
Summary of clinical features, immune escape mechanisms and subtype-specific genomic alterations of three TMEC subtypes.

The potential immune escape mechanism of each subtype may have important clinical implications. The immune-active subtype has increased immune-suppressive cells, high immune-suppressive cytokine levels, and upregulating immune inhibitors, implying both intense tumour-suppressing and tumour-promoting immune responses existing. Theoretically, such subtype may benefit most from immune checkpoint inhibitors (ICI), the most studied immunotherapy targeting TME in CRC, and MSI status is a solid predictive marker for ICI efficacy[40]. However, not all dMMR/MSI-H patients respond to anti-PD1/anti-PD-L1 therapy. Previous researches reported an ORR of 31% in refractory mCRC[31] and 44% in the first-line setting[41]. Most dMMR/MSI-H CRC is of immune-active subtype, targeting the potential immune escape mechanism may be a novel strategy to improve the efficacy. Strategies such as erasing the infiltration of immunosuppressive cells, neutralising immunosuppressive cytokines, and combining other checkpoint inhibitors can be considered to activate tumour immunity further.

The immune-desert subtype has decreased immune-promotive cells, low immune-promotive cytokine level, downregulating immune inhibitors, and defective antigen presentation. Nevertheless, the OS is similar to the immune-active subtype, implying the survival impairment of stromal cells has neutralised the survival benefit of high infiltration of activated adaptive immune cells. Even the infiltration of immunosuppressive cells, such as MDSC and Treg, may improve prognosis, which shows the complexity and heterogeneity of CRC TME. The major difference between the immune-desert subtype (“cold tumour”) to other two subtypes (“hot tumour”) is the antigen presentation defect. Therefore, increasing tumour immunogenicity may induce the immune cell chemotaxis and transform a “cold tumour” into a “hot tumour”. Chemotherapy and radiotherapy can cause tumour cell death with subsequent release of cellular fragmentation and cancer-associated neoantigens, which are presented to APCs to increase tumour immunogenicity[42]. The use of pembrolizumab plus FOLFOX achieved ORR 55% in the first-line setting[43]. However, the synergistic effect of pembrolizumab and radiotherapy was not as expected, which only achieved an ORR of 4.5% in 22 pMMR/MSS mCRC patients[44]. Further researches of a dual checkpoint inhibitor (anti-PD-L1 + anti-CTLA-4) following radiotherapy are underway (NCT02701400, NCT03122509). Other novel approaches like tumour vaccine, and oncolytic virus are under early-phase research.

The stroma-rich subtype, which takes up nearly half the samples, has a similar immune escape mechanism with the immune-active subtype but decreased immune-promotive cells. This subtype is “hot” but owns the worst survival. The primary reason may be the pro-tumoural effect of excessive stromal cell infiltration, excluding activated adaptive immune cells from the tumour. Anti-fibroblast, anti-TGF-β pathway, anti-angiogenesis, anti-immunosuppressive cytokines, or the combination of the above therapy may be hoping to transform it into an immune-active subtype. Dual antagonising of TGF-β and PD-1/PD-L1 showed promising results in preclinical researches[45]. The combination of vactosertib (a small-molecule inhibitor of TGF-β) and pembrolizumab showed an ORR of 15.2% in previously treated MSS mCRC patients[46]. Furthermore, REGONIVO (regorafenib + nivolumab) trial reported inspiring 36% ORR in unselected mCRC patients[47], which implies the potential of ICI and anti-angiogenesis. Previous studies showed that cancer-associated fibroblasts (CAFs) could secret tumourigenic factors, modulate immunosuppressive TME, and create an ECM barrier to block CD8 T cells accessing the tumour[48]. Multiple strategies targeting CAFs are under research, such as normalising the phenotype of CAFs, inhibiting CAF generation and activation, reducing CAF secretome[49], et al., and awaiting the results.

Also, the limitation of this study cannot be ignored. The major one is that the subtyping was based on transcriptomic data, which needed to be further validated pathologically. Moreover, the immune escape mechanism speculation was mainly based on in silico analysis, and numerous results only reflected the correlation but not the causation. Further in vitro and in vivo experiments were needed.

In conclusion, we systematically presented the infiltration pattern of CRC TME cells. CRC samples can be divided into three subtypes with distinct TME cell infiltration patterns, clinical features, genomic characteristics, as well as their immune escape mechanism underneath. The results may provide inspiration and direction for further researches on CRC immunotherapy.

## 5 Declarations

### 5.1 Ethics approval and consent to participate

Not applicable.

### 5.2 Consent for publication

All authors gave consent for publication.

### 5.3 Availability of data and materials

All datasets mentioned in this article were publicly available.

### 5.6 Authors’ contributions

YM, GH and JX contributed to the planning of the study. YM, QF and YX collected the data and performed most of the bioinformatics analysis. YL and WC verified the numerical results by an independent implementation. YM, QF, YX, and YL drafted and revised the manuscript. All authors contributed to the interpretation of data and review of the manuscript. All authors reviewed and approved the final manuscript.

## 5.7 Acknowledgements

Not applicable.

