## Supplementary material for "The infiltration pattern of microenvironmental cells and different immune escape mechanisms in colorectal cancer": Fig. S

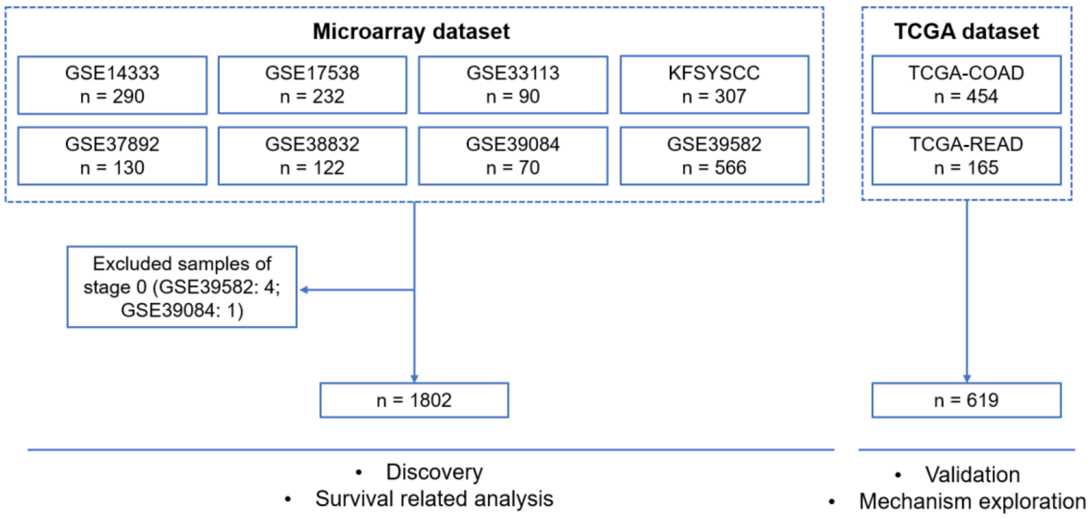


Figure S1. The composition of microarray and TCGA datasets.

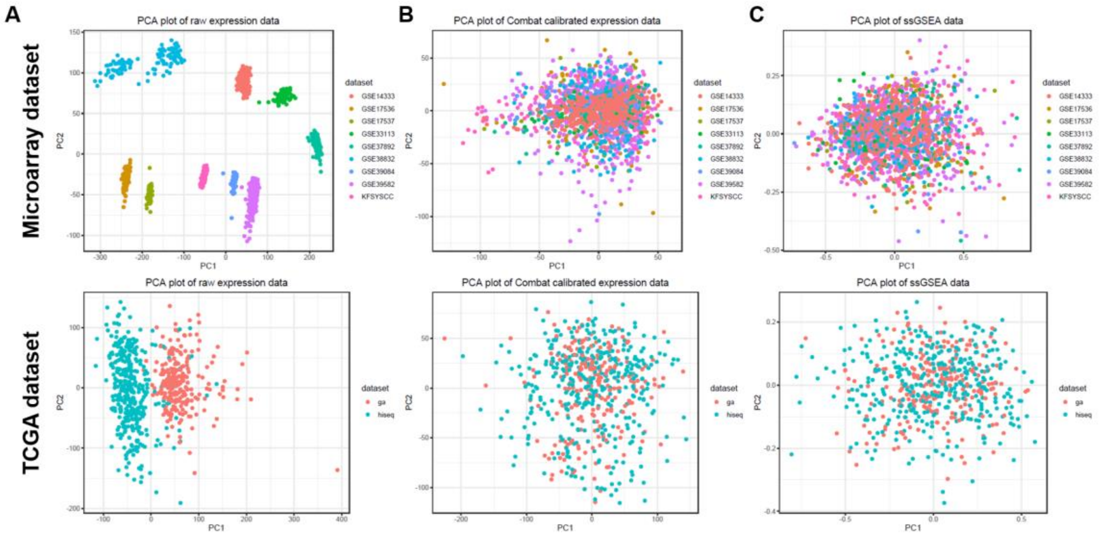


Figure S2. PCA plots of expression data before (A) and after (B) Combat algorithm calibration for batch effect and ssGSEA results based on calibrated expression data (C).

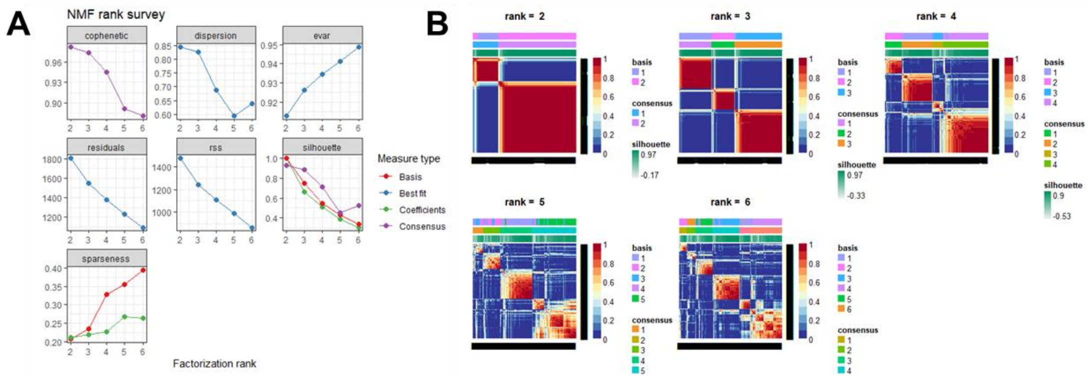


Figure S3. The NMF rank survey (A) and heatmaps of NMF factors (B) in the microarray dataset.

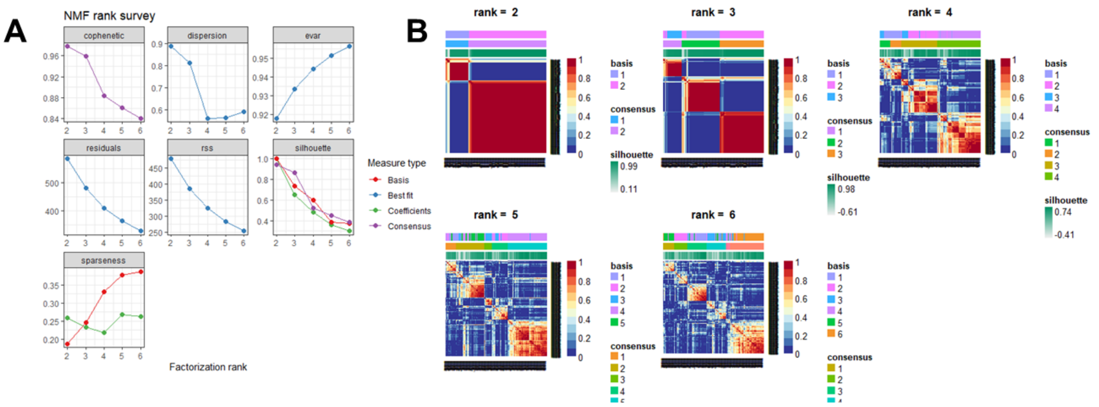

Figure S4. The NMF rank survey (A) and heatmaps of NMF factors (B) in the TCGA dataset.

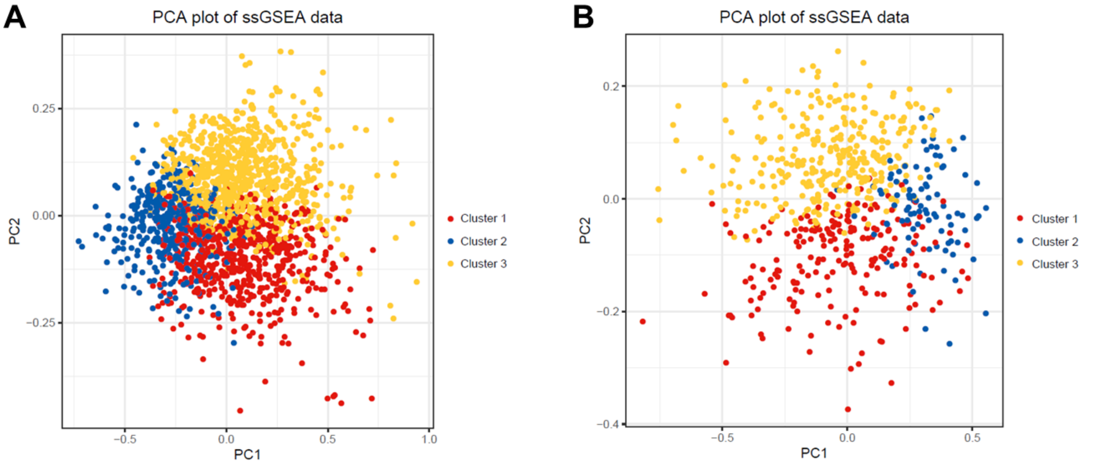

Figure S5. The PCA plots of ssGSEA results annotated with classification results of microarray (A) and TCGA (B) dataset.

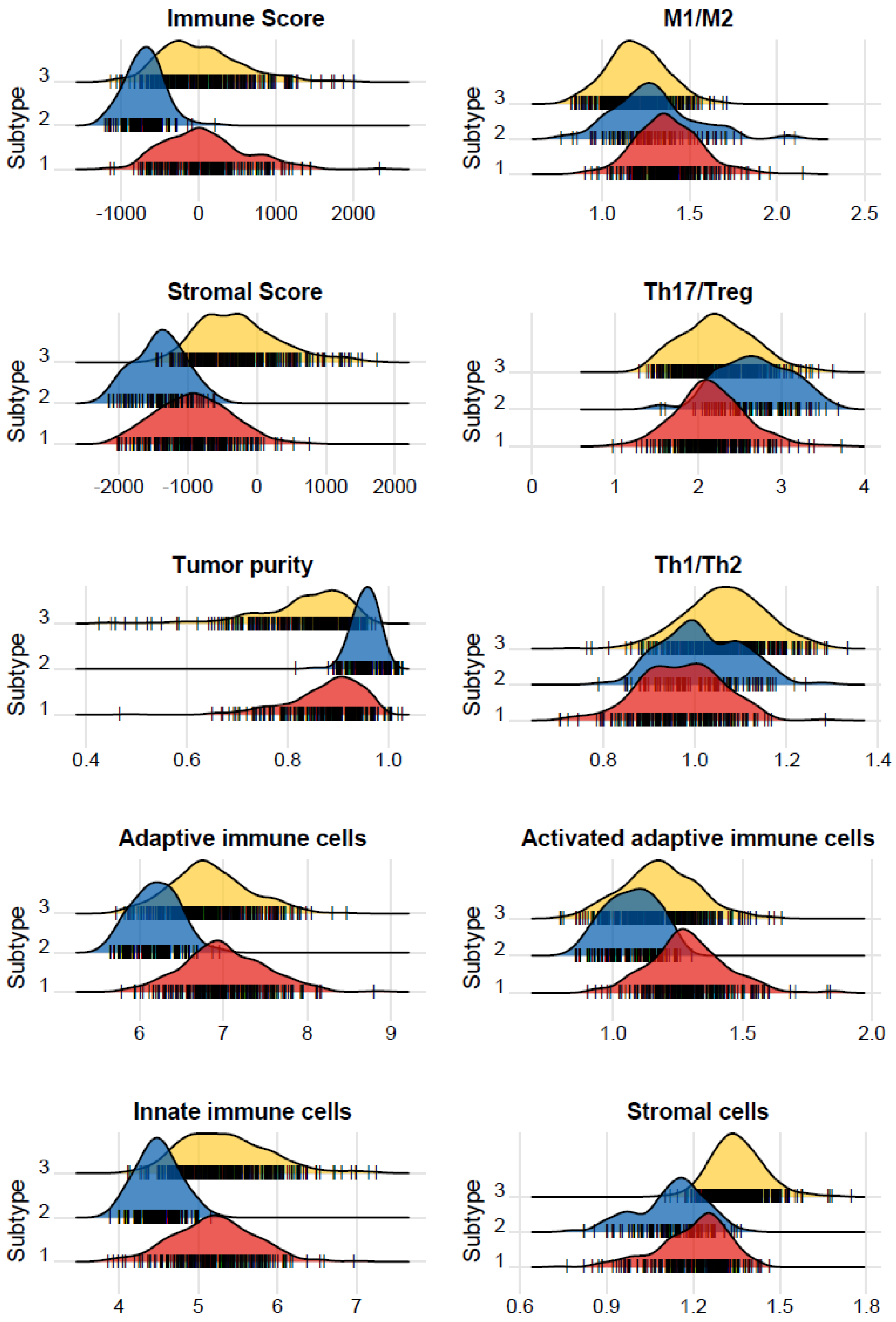


Figure S6. The ESTIMATE results and other characteristics of the TCGA dataset.

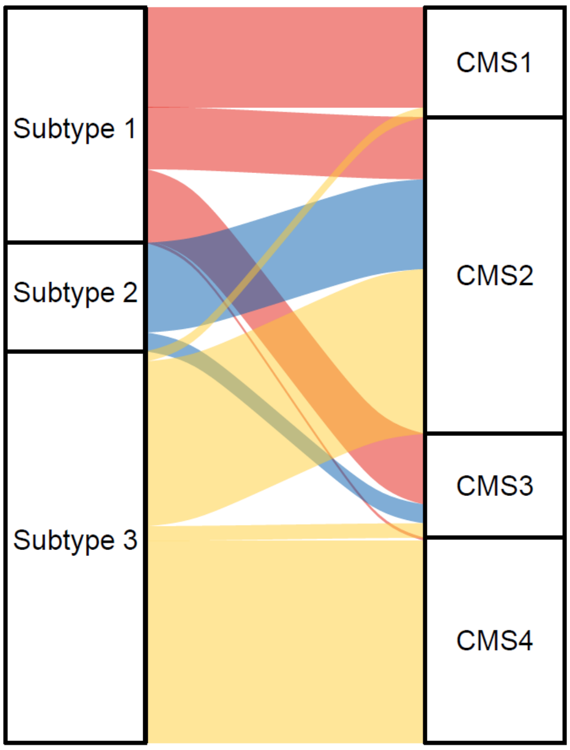


Figure S7. The Sankey plot comparing the distribution of TMECS and CMS in TCGA dataset.

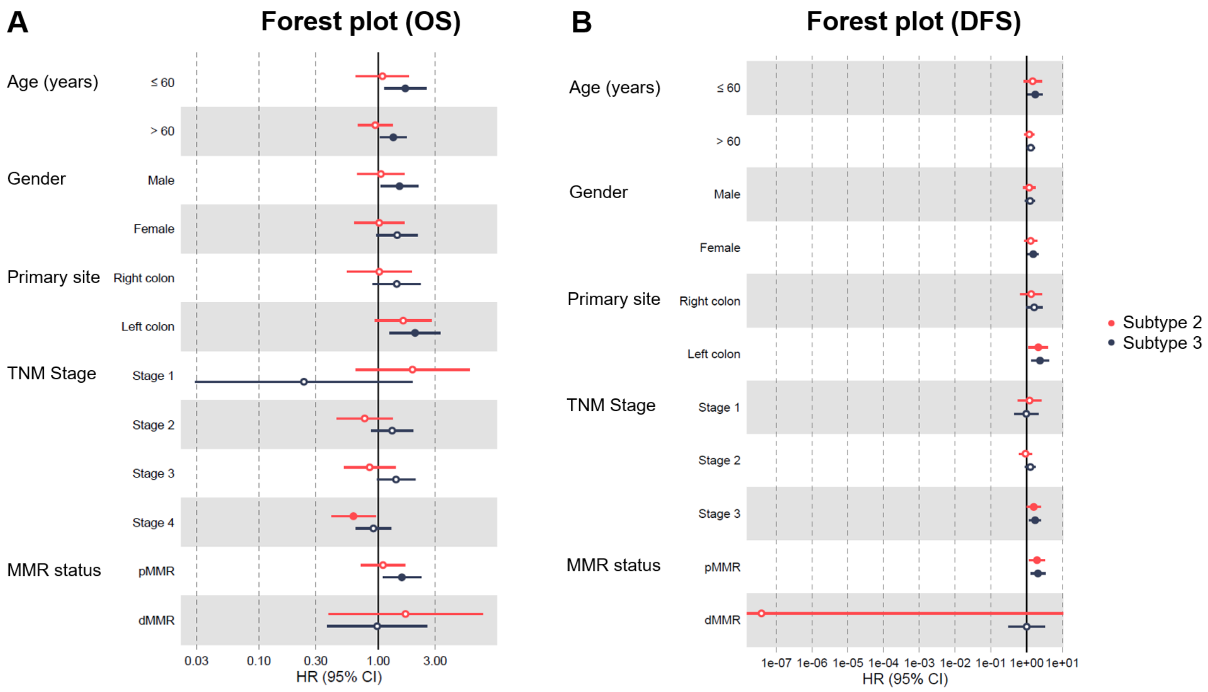


Figure S8. Subgroup analyses of OS (A) and DFS (B) in microarray dataset. Factors with P value less than 0.05 were drawn with solid points.
