## Supplementary material for "The infiltration pattern of microenvironmental cells and different immune escape mechanisms in colorectal cancer": Table S

Table S1. The detailed gene signature used in our ssGSEA algorithm

| **Cell types** | **Origin** | **Gene signature** |
| --- | --- | --- |
| M0 macrophage | Xiao Y et al. | BHLHE41;CHI3L1;COL8A2;CSF1;CXCL5;CYP27A1;DCSTAMP;GPC4;MARCO;MMP9;PLA2G7;PPBP |
| M1 macrophage | Xiao Y et al. | ACHE;ADAMDEC1;APOL3;APOL6;ARRB1;CCL19;CCL8;CD40;CXCL10;CXCL11;CXCL9;CYP27B1;EBI3;HESX1;MACF1;NOD2;PLA1A;SIGLEC1;SLAMF1;SLC2A6;SOCS1;TLR7;TNFAIP6;TNIP3 |
| M2 macrophage | Xiao Y et al. | CCL14;CCL18;CCL23;CD4;CD68;CLEC4A;CRYBB1;FRMD4A;HRH1;MS4A6A;NME8;NPL;RENBP;WNT5B |
| Endothelial cell | MCPcounter | ACVRL1;APLN;BCL6B;BMP6;BMX;CDH5;CLEC14A;CXorf36;EDN1;ELTD1;EMCN;ESAM;ESM1;HECW2;HHIP;KDR;MMRN1;MMRN2;MYCT1;PALMD;PEAR1;PGF;PLXNA2;PTPRB;ROBO4;SDPR;SHANK3;SHE;TEK;TIE1;VEPH1;VWF |
| Fibroblast | MCPcounter | COL1A1;COL3A1;COL6A1;COL6A2;DCN;GREM1;PAMR1;TAGLN |
| Activated B cell | CHAROENTONG P et al. | ADAM28;CD180;CD79B;BLK;CD19;MS4A1;TNFRSF17;IGHM;GNG7;MICAL3;SPIB;HLA-DOB;IGKC;PNOC;FCRL2;BACH2;CR2;TCL1A;AKNA;ARHGAP25;CCL21;CD27;CD38;CLEC17A;CLEC9A;CLECL1 |
| Activated CD4 T cell | CHAROENTONG P et al. | AIM2;BIRC3;BRIP1;CCL20;CCL4;CCL5;CCNB1;CCR7;DUSP2;ESCO2;ETS1;EXO1;EXOC6;IARS;ITK;KIF11;KNTC1;NUF2;PRC1;PSAT1;RGS1;RTKN2;SAMSN1;SELL;TRAT1 |
| Activated CD8 T cell | CHAROENTONG P et al. | ADRM1;AHSA1;C1GALT1C1;CCT6B;CD37;CD3D;CD3E;CD3G;CD69;CD8A;CETN3;CSE1L;GEMIN6;GNLY;GPT2;GZMA;GZMH;GZMK;IL2RB;LCK;MPZL1;NKG7;PIK3IP1;PTRH2;TIMM13;ZAP70 |
| Central memory CD4 T cell | CHAROENTONG P et al. | ABHD3;AHNAK;ANXA2P2;AQP3;ATHL1;BMI1;BZW2;CD63;COL4A1;CYLD;ELMO2;FYN;GLIPR1;GSS;IFITM2;ITGB1;ITGB2;KLF5;LSP1;NDUFB9;PKM2;SFXN3;SIRPG;SMAD4;STX4;TRADD;VIM;XRCC6 |
| Central memory CD8 T cell | CHAROENTONG P et al. | ACTN4;ADAM12;ADCY9;F13A1;FCER1G;FCGR3B;FGF7;FKBP4;GLUD1;GM2A;GUSB;IL1RN;NOL11;NTRK1;RARA;RNF128;SIGLEC1;TNFRSF11A;TOX4;UBA52;ULBP1 |
| Effector memeory CD4 T cell | CHAROENTONG P et al. | ATM;CASP3;CASQ1;CD300E;DARS;DOCK9;EXOSC9;EZH2;GDE1;IL34;NCOA4;NEFL;PDGFRL;PTGS1;REPS1;SCG2;SDPR;SIGLEC14;SIGLEC6;TAL1;TFEC;TIPIN;TPK1;UQCRB;USP9Y;WIPF1;ZCRB1 |
| Effector memeory CD8 T cell | CHAROENTONG P et al. | ACAP1;APOL3;ARHGAP10;ATP10D;C3AR1;CCR5;CD160;CD55;CFLAR;CMKLR1;DAPP1;FCRL6;FLT3LG;GZMM;HAPLN3;HLA-DMB;HLA-DPA1;HLA-DPB1;IFI16;LIME1;LTK;NFKBIA;SETD7;SIK1;TRIB2 |
| Gamma delta T cell | CHAROENTONG P et al. | ACP5;AQP9;BTN3A2;C1orf54;CARD8;CCL18;CD209;CD33;CD36;CDK5;IL10RB;KLRF1;LGALS1;MAPK7;KLHL7;KRT80;LAMC1;LCORL;LMNB1;MEIS3P1;MPL;FABP1;FABP5;FADD;MFAP3L;MINPP1;RPS24;RPS7;RPS9;DBNL;CCL13 |
| Immature B cell | CHAROENTONG P et al. | CD22;CYBB;FAM129C;FCRL1;FCRL3;FCRL5;FCRLA;HDAC9;HLA-DQA1;HVCN1;KIAA0226;NCF1;NCF1B;P2RY10;SP100;TXNIP;STAP1;TAGAP;ZCCHC2 |
| Memory B cell | CHAROENTONG P et al. | AICDA;CCNA2;CDKN3;CLCN5;ENPP1;FCER1A;FCRL4;MYC;RUNX2;SORL1;SOX5;STAT5A;STAT5B;TLR9 |
| Regulatory T cell | Xiao Y et al. | BARX2;CD5;CD7;CD70;CEMP1;CRISP3;CTLA4;EFNA5;FOXP3;FRMD8;HIC1;HMGB3P30;KIRREL;LAIR2;LILRA4;LOC126987;NPAS1;NTN3;PLCH2;PMCH;RYR1;SEC31B;KLHL22;SIT1;SKAP1;SSX1;TRAV21;TRAV8-6;TRAV9-2 |
| T follicular helper cell | CHAROENTONG P et al. | B3GAT1;CDK5R1;PDCD1;BCL6;CD200;CD83;CD84;FGF2;GPR18;CEBPA;CECR1;CLEC10A;CLEC4A;CSF1R;CTSS;DMN;DPP4;LRRC32;MC5R;MICA;NCAM1;NCR2;NRP1;PDCD1LG2;PDCD6;PRDX1;RAE1;RAET1E;SIGLEC7;SIGLEC9;TYRO3;CHST12;CLIC3;IVNS1ABP;KIR2DL2;LGMN |
| Type 1 T helper cell | CHAROENTONG P et al. | CD70;TBX21;ADAM8;AHCYL2;ALCAM;B3GALNT1;BBS12;BST1;CD151;CD47;CD48;CD52;CD53;CD59;CD6;CD68;CD7;CD96;CFHR3;CHRM3;CLEC7A;COL23A1;COL4A4;COL5A3;DAB1;DLEU7;DOC2B;EMP1;F12;FURIN;GAB3;GATM;GFPT2;GPR25;GREM2;HAVCR1;HSD11B1;HUNK;IGF2;RCSD1;RYR1;SAV1;SELE;SELP;SH3KBP1;SIT1;SLC35B3;SIGLEC10;SKAP1;THUMPD2;TIGIT;ZEB2;ENC1;FAM134B;FBXO30;FCGR2C;STAC;LTC4S;MAN1B1;MDH1;MMD;RGS16;IL12A;P2RX5;CD97;ITGB4;ICAM3;METRNL;TNFRSF1A;IRF1;HTR2B;CALD1;MOCOS;TRAF3IP2;TLR8;TRAF1;DUSP14 |
| Type 17 T helper cell | CHAROENTONG P et al. | IL17A;IL17RA;C2CD4A;C2CD4B;CA2;CCDC65;CEACAM3;IL17C;IL17F;IL17RC;IL17RE;IL23A;ILDR1;LONRF3;SH2D6;TNIP2;ABCA1;ABCB1;ADAMTS12;ANK1;ANKRD22;B3GALT2;CAMTA1;CCR9;CD40;GPR44;IFT80 |
| Type 2 T helper cell | CHAROENTONG P et al. | ASB2;CSRP2;DAPK1;DLC1;DNAJC12;DUSP6;GNAI1;LAMP3;NRP2;OSBPL1A;PDE4B;PHLDA1;PLA2G4A;RAB27B;RBMS3;RNF125;TMPRSS3;GATA3;BIRC5;CDC25C;CDC7;CENPF;CXCR6;DHFR;EVI5;GSTA4;HELLS;IL26;LAIR2 |
| Activated dendritic cell | Xiao Y et al. | ARHGAP22;BIRC3;CCL17;CCL22;CD86;CHST7;CLIC2;ETV3;HTR2B;IL12B;MAP3K13;PDCD1LG2 |
| Resting natural killer cell | Xiao Y et al. | CDHR1;DEFA4;KLRC3;KLRF1;NAALADL1;S1PR5;TEP1;TTC38;ZNF135 |
| Activated natural killer cell | Xiao Y et al. | CCND2;CDK6;CTSW;GZMA;IL12RB2;KIR2DL1;KIR2DL4;KIR2DS4;KIR3DL2;NCR3;TNFSF14 |
| Eosinophil | CHAROENTONG P et al. | GIPR;KRT18P50;LRMP;FOSB;RRP12;GPR183;NR4A3;ST3GAL6;DEPDC5;PDE6C;PKD2L2;GPR65;IL5RA;P2RY14;DACH1;DAPK2;EMR3 |
| Resting dendritic cell | Xiao Y et al. | ALOX15;C1orf54;CD1A;CD1B;CD1C;CD1E;DHRS11;MMP12;PPFIBP1;RNASE6;SCN9A;TREM2 |
| Mast cell | CHAROENTONG P et al. | ADAMTS3;CPA3;CMA1;CTSG;ARHGAP15;CPM;FCN1;FTL;HSPA6;ITGA9;RNASE3;S100A4;SIGLEC8;SLC6A4;PTGS2;EGR3;PILRA |
| MDSC | CHAROENTONG P et al. | CCR2;CD14;CD2;CD86;CXCR4;FCGR2A;FCGR2B;FCGR3A;FERMT3;GPSM3;IL18BP;IL4R;ITGAL;ITGAM;PARVG;PSAP;PTGER2;PTGES2;S100A8;S100A9 |
| Monocyte | CHAROENTONG P et al. | ASGR2;CFP;ASGR1;CD1D;UPK3A;ACTG1;ANXA5;ATP6V1B2;CFL1;DAZAP2;CTBS;EMR4P;HIVEP2;MARCKSL1;MBP;MMP15;PNPLA6;TMBIM6;PQBP1;TEX264;IKZF1 |
| Natural killer T cell | CHAROENTONG P et al. | BTN2A2;CD101;CD109;CNPY3;CNPY4;CREB1;CRTC2;CRTC3;CSF2;KLRC1;FUT4;ICAM2;IL32;LAMP2;LILRB5;KLRG1;HSPA4;HSPB6;ISM2;ITIH2;KDM4C;KIR2DS4;KIRREL3;SDCBP;NFATC2IP;MICB;KIR2DL1;KIR2DL3;KIR3DL1;KIR3DL2;NCR1;FOSL1;TSLP;SLC7A7;SPP1;TREM2;UBASH3A;YBX2;CCDC88A;CLEC1A;THBD;PDPN;VCAM1;EMR1 |
| Neutrophil | CHAROENTONG P et al. | CREB5;CDA;CHST15;S100A12;APOBEC3A;CASP5;MMP25;HAL;C1orf183;FFAR2;MAK;CXCR1;STEAP4;MGAM;BTNL8;CXCR2;TNFRSF10C;VNN3 |
| Plasmacytoid dendritic cell | CHAROENTONG P et al. | CBX6;DAB2;DDX17;HIGD1A;IDH3A;IL3RA;MAGED1;NUCB2;OFD1;OGT;PDIA4;SERTAD2;SIRPA;TMED2;ENG;FCAR;IGF1;ITGA2B;GABARAP;GPX1;KRT23;PROK2;RALB;RETNLB;RNF141;SEC14L1;SEPX1;EMP3;CD300LF;ABTB1;KLHL21;PHRF1 |

Table S2. Correlation between our ssGSEA method and CIBERSORT or MCP-counter in microarray dataset

| **Origin** | **Cell names in CIBERSORT or MCP-counter** | **Cell names in our method** | **Correlation (R)** | **P value** |
| --- | --- | --- | --- | --- |
| CIBERSORT | B.cells.naive | Immature B cell | 0.4975377 | < 2.2e-16 |
| CIBERSORT | B.cells.memory | Memory B cell | 0.09038173 | 0.0001221 |
| CIBERSORT | Plasma.cells | Activated B cell | 0.6353449 | < 2.2e-16 |
| CIBERSORT | T.cells.CD8 | Activated CD8 T cell + Central memory CD8 T cell + Effector memeory CD8 T cell | 0.5518634 | < 2.2e-16 |
| CIBERSORT | T.cells.CD4.naive + T.cells.CD4.memory.resting + T.cells.CD4.memory.activated | Central memory CD4 T cell + Effector memeory CD4 T cell + Activated CD4 T cell | 0.157468 | 1.79E-11 |
| CIBERSORT | T.cells.follicular.helper | T follicular helper cell | 0.339686 | < 2.2e-16 |
| CIBERSORT | T.cells.regulatory..Tregs. | Regulatory T cell | 0.1976455 | < 2.2e-16 |
| CIBERSORT | T.cells.gamma.delta | Gamma delta T cell | 0.338848 | < 2.2e-16 |
| CIBERSORT | NK.cells.resting | Resting natural killer cell | 0.1320923 | 1.82E-08 |
| CIBERSORT | NK.cells.activated | Activated natural killer cell | 0.1992691 | < 2.2e-16 |
| CIBERSORT | Monocytes | Monocyte | 0.144679 | 6.84E-10 |
| CIBERSORT | Macrophages.M0 | M0 macrophage | 0.6640603 | < 2.2e-16 |
| CIBERSORT | Macrophages.M1 | M1 macrophage | 0.7698678 | < 2.2e-16 |
| CIBERSORT | Macrophages.M2 | M2 macrophage | 0.6707435 | < 2.2e-16 |
| CIBERSORT | Dendritic.cells.resting | Resting dendritic cell | 0.4073389 | < 2.2e-16 |
| CIBERSORT | Dendritic.cells.activated | Activated dendritic cell | 0.1992691 | < 2.2e-16 |
| CIBERSORT | Mast.cells.resting + Mast.cells.activated | Mast cell | 0.3836081 | < 2.2e-16 |
| CIBERSORT | Eosinophils | Eosinophils | 0.3077271 | < 2.2e-16 |
| CIBERSORT | Neutrophils | Neutrophil | 0.636979 | < 2.2e-16 |
| MCP-counter | Endothelial cells | Endothelial cell | 0.9689445 | < 2.2e-16 |
| MCP-counter | Fibroblasts | Fibroblast | 0.9696201 | < 2.2e-16 |

Table S3. Correlation between our ssGSEA method and CIBERSORT or MCP-counter in TCGA dataset

| **Origin** | **Cell names in CIBERSORT or MCP-counter** | **Cell names in our method** | **Correlation (R)** | **P value** |
| --- | --- | --- | --- | --- |
| CIBERSORT | B.cells.naive | Immature B cell | 0.3459307 | < 2.2e-16 |
| CIBERSORT | B.cells.memory | Memory B cell | -0.01252574 | 0.7558 |
| CIBERSORT | Plasma.cells | Activated B cell | 0.2409421 | 1.26E-09 |
| CIBERSORT | T.cells.CD8 | Activated CD8 T cell + Central memory CD8 T cell + Effector memeory CD8 T cell | 0.3334561 | < 2.2e-16 |
| CIBERSORT | T.cells.CD4.naive + T.cells.CD4.memory.resting + T.cells.CD4.memory.activated | Central memory CD4 T cell + Effector memeory CD4 T cell + Activated CD4 T cell | 0.3879787 | < 2.2e-16 |
| CIBERSORT | T.cells.follicular.helper | T follicular helper cell | 0.2166525 | 5.21E-08 |
| CIBERSORT | T.cells.regulatory..Tregs. | Regulatory T cell | 0.4264065 | < 2.2e-16 |
| CIBERSORT | T.cells.gamma.delta | Gamma delta T cell | NA | NA |
| CIBERSORT | NK.cells.resting | Resting natural killer cell | 0.0124071 | 7.58E-01 |
| CIBERSORT | NK.cells.activated | Activated natural killer cell | 0.3606555 | < 2.2e-16 |
| CIBERSORT | Monocytes | Monocyte | 0.03423788 | 3.95E-01 |
| CIBERSORT | Macrophages.M0 | M0 macrophage | 0.6510624 | < 2.2e-16 |
| CIBERSORT | Macrophages.M1 | M1 macrophage | 0.7502583 | < 2.2e-16 |
| CIBERSORT | Macrophages.M2 | M2 macrophage | 0.6581065 | < 2.2e-16 |
| CIBERSORT | Dendritic.cells.resting | Resting dendritic cell | 0.2045936 | 2.83E-07 |
| CIBERSORT | Dendritic.cells.activated | Activated dendritic cell | 0.1306583 | 1.12E-03 |
| CIBERSORT | Mast.cells.resting + Mast.cells.activated | Mast cell | 0.09480138 | 0.01832 |
| CIBERSORT | Eosinophils | Eosinophils | 0.17368 | 1.39E-05 |
| CIBERSORT | Neutrophils | Neutrophil | 0.5539531 | < 2.2e-16 |
| MCP-counter | Endothelial cells | Endothelial cell | 0.904981 | < 2.2e-17 |
| MCP-counter | Fibroblasts | Fibroblast | 0.9479525 | < 2.2e-18 |

Table S4. The association between TME cell subtypes and clinicopathological features in the microarray dataset

|  | **Cluster 1** | **Cluster 2** | **Cluster 3** | **P value** |
| --- | --- | --- | --- | --- |
| **N** | 594 | 404 | 804 |  |
| **Age (mean±SD)** | 66.83±14.48 | 65.87±13.70 | 64.22±12.99 | **0.005** |
| **Gender** |  |  |  | **<0.001** |
| **Male** | 190 (32.0) | 153 (37.9) | 327 (40.7) |  |
| **Female** | 217 (36.5) | 131 (32.4) | 226 (28.1) |  |
| **Unknown** | 187 (31.5) | 120 (29.7) | 251 (31.2) |  |
| **TNM stage** |  |  |  | **<0.001** |
| **1** | 87 (14.6) | 34 (8.4) | 43 (5.3) |  |
| **2** | 242 (40.7) | 175 (43.3) | 318 (39.6) |  |
| **3** | 194 (32.7) | 123 (30.4) | 286 (35.6) |  |
| **4** | 71 (12.0) | 72 (17.8) | 157 (19.5) |  |
| **Site** |  |  |  | **<0.001** |
| **Right colon** | 153 (25.8) | 61 (15.1) | 144 (17.9) |  |
| **Left colon** | 125 (21.0) | 135 (33.4) | 224 (27.9) |  |
| **Rectum** | 1 (0.2) | 0 (0.0) | 7 (0.9) |  |
| **Unknown** | 315 (53.0) | 208 (51.5) | 429 (53.4) |  |
| **MMR status** |  |  |  | **<0.001** |
| **dMMR** | 57 (9.6) | 4 (1.0) | 29 (3.6) |  |
| **pMMR** | 135 (22.7) | 137 (33.9) | 223 (27.7) |  |
| **Unknown** | 402 (67.7) | 263 (65.1) | 552 (68.7) |  |
| **CMS subtype** |  |  |  | **<0.001** |
| **CMS1** | 147 (24.7) | 4 (1.0) | 54 (6.7) |  |
| **CMS2** | 133 (22.4) | 240 (59.4) | 172 (21.4) |  |
| **CMS3** | 95 (16.0) | 69 (17.1) | 22 (2.7) |  |
| **CMS4** | 34 (5.7) | 2 (0.5) | 311 (38.7) |  |
| **NOLBL** | 53 (8.9) | 22 (5.4) | 65 (8.1) |  |
| **Unknown** | 132 (22.2) | 67 (16.6) | 180 (22.4) |  |

Note: bold values present p values where p < 0.05. Abbreviations: SD: standard deviation; TNM: tumor node metastasis; MMR: DNA mismatch repair; CMS: Consensus Molecular Subgroups; NOLBL: samples that could not be attributed to any CMS subtype

Table S5. The association between TME cell subtypes and clinicopathological features in the TCGA dataset

|  | **Cluster 1** | **Cluster 2** | **Cluster 3** | **P value** |
| --- | --- | --- | --- | --- |
| **N** | 203 | 92 | 324 |  |
| **Age (mean±SD)** | 68.37±13.51 | 65.25±10.98 | 65.31±12.63 | **0.019** |
| **Gender** |  |  |  | 0.669 |
| **Male** | 107 (52.7) | 53 (57.6) | 170 (52.5) |  |
| **Female** | 96 (47.3) | 39 (42.4) | 154 (47.5) |  |
| **TNM stage** |  |  |  | **0.001** |
| **1** | 42 (20.7) | 16 (17.4) | 48 (14.8) |  |
| **2** | 93 (45.8) | 34 (37.0) | 102 (31.5) |  |
| **3** | 50 (24.6) | 24 (26.1) | 108 (33.3) |  |
| **4** | 13 (6.4) | 17 (18.5) | 59 (18.2) |  |
| **Unknown** | 5 (2.5) | 1 (1.1) | 7 (2.2) |  |
| **Site** |  |  |  | **<0.001** |
| **Right colon** | 129 (63.5) | 28 (30.4) | 99 (30.6) |  |
| **Left colon** | 32 (15.8) | 37 (40.2) | 113 (34.9) |  |
| **Rectum** | 34 (16.7) | 25 (27.2) | 101 (31.2) |  |
| **Unknown** | 8 (3.9) | 2 (2.2) | 11 (3.4) |  |
| **MSI status** |  |  |  | **<0.001** |
| **MSI-H** | 69 (34.0) | 0 (0.0) | 13 (4.0) |  |
| **MSI-L** | 25 (12.3) | 15 (16.3) | 53 (16.4) |  |
| **MSS** | 97 (47.8) | 64 (69.6) | 243 (75.0) |  |
| **Unknown** | 12 (5.9) | 13 (14.1) | 15 (4.6) |  |
| **CMS subtype** |  |  |  | **<0.001** |
| **CMS1** | 69 (34.0) | 0 (0.0) | 7 (2.2) |  |
| **CMS2** | 43 (21.2) | 62 (67.4) | 114 (35.2) |  |
| **CMS3** | 49 (24.1) | 13 (14.1) | 10 (3.1) |  |
| **CMS4** | 2 (1.0) | 0 (0.0) | 140 (43.2) |  |
| **NOLBL** | 25 (12.3) | 2 (2.2) | 33 (10.2) |  |
| **Unknown** | 15 (7.4) | 15 (16.3) | 20 (6.2) |  |

Note: bold values present p values where p < 0.05. Abbreviations: SD: standard deviation; TNM: tumor node metastasis; MSI: microsatellite instability; CMS: Consensus Molecular Subgroups; NOLBL: samples that could not be attributed to any CMS subtype

Table S6. Univariate and multivariate Cox regression analysis for OS in microarray dataset

| **Factors** | **Univariate analysis** |  | **Multivariate analysis** |  |
| --- | --- | --- | --- | --- |
|  | **HR (95% CI)** | ***P*** | **HR (95% CI)** | ***P*** |
| **Age (years)** |  |  |  |  |
| **≤ 60** | 1 (reference) |  |  |  |
| **> 60** | 1.038 (0.852 - 1.265) | 0.709 |  |  |
| **Gender** |  |  |  |  |
| **Male** | 1 (reference) |  |  |  |
| **Female** | 0.827 (0.660 - 1.037) | 0.1 |  |  |
| **Primary site** |  |  |  |  |
| **Right-sided colon** | 1 (reference) |  |  |  |
| **Left-sided colon** | 0.927 (0.704 - 1.221) | 0.592 |  |  |
| **Rectum** | 2.000 (0.379 – 3.794) | 0.757 |  |  |
| **TNM Stage** |  |  |  |  |
| **I** | 1 (reference) |  | 1 (reference) |  |
| **II** | 2.092 (1.202 – 3.643) | **0.009** | 2.069 (1.187 – 3.605) | **0.01** |
| **III** | 3.134 (1.813 – 5.419) | **<0.001** | 3.070 (1.773 – 5.316) | **<0.001** |
| **IV** | 14.434 (8.370 – 24.891) | **<0.001** | 14.249 (8.236 – 24.650) | **<0.001** |
| **MMR status** |  |  |  |  |
| **MMR-proficient** | 1 (reference) |  |  |  |
| **MMR-deficient** | 0.732 (0.470 – 1.141) | 0.169 |  |  |
| **TME cell subtype** |  |  |  |  |
| **Cluster 1** | 1 (reference) |  |  |  |
| **Cluster 2** | 0.968 (0.741 – 1.264) | 0.81 | 0.770 (0.589 – 1.008) | 0.057 |
| **Cluster 3** | 1.449 (1.174 – 1.787) | **<0.001** | 1.151 (0.931 – 1.423) | 0.195 |

Note: bold values present p values where p < 0.05. Abbreviations: SD: standard deviation; TNM: tumor node metastasis; MMR: DNA mismatch repair; CMS: Consensus Molecular Subgroups; HR: hazard ratio
